# Lifetime changes in Body map based on music prediction

**DOI:** 10.1101/2025.02.02.636100

**Authors:** Masaki Tanaka, Tatsuya Daikoku

## Abstract

This study aims to investigate the relationship between music, which is said to promote well-being, and emotions as well as bodily sensations across the lifespan. Using eight chord progressions based on predictive models, we conducted online experiments with 784 participants aged 10 to 80, evaluating bodily sensation mapping and emotional responses. The results revealed that bodily sensations peaked in the 20s to 30s and subsequently declined with age, particularly for chords with low prediction error and uncertainty. On the other hand, arousal levels increased with age and showed a strong correlation with predictable musical structures. These findings suggest that bottom-up bodily responses diminish with ageing, while supporting the relationship between top-down processing shaped by accumulated experience and the predictability of music, as indicated by previous research. This study enhances the development of music-based interventions that improve well-being across generations, considering the unique physical and emotional characteristics of each age group.

## Introduction

Music has often been recognized as a medium through which individuals can express and embody their emotional experiences, subsequently fostering their overall well-being (Juslin & Sloboda, 2001). Well-being, in turn, is associated with a reduction in the risks of illness and mortality (Trudel-Fitzgerald et al., 2019) and tends to correlate with higher well-being scores among healthy individuals (Dirzyte & Perminas, 2021)(for review, see Hallam and Himonides, 2022). In light of the pervasive stagnation experienced globally following the COVID-19 pandemic, the importance of well-being has received heightened attention across various sectors worldwide.

Previous research has shown that music is linked to various psychological elements and plays a significant role in maintaining and altering states such as mood, emotion, arousal levels, and reminiscence (Juslin & Laukka, 2004; Schafer et al., 2013; Hallam and Himonides, 2022). Music is appreciated across all age groups, from children to the elderly and adults alike. Sloboda and colleagues (2009) found that music listening in adults’ daily lives significantly influences emotions, including reminiscence and mood. In younger individuals, music has been studied as a tool for emotional regulation and cognitive adjustment (Zoe et al., 2015). Additionally, Laukka (2007) discovered that music listening among older adulthood frequently elicits positive emotions, which contributes to their psychological well-being.

Numerous studies have demonstrated that music significantly affects bodily sensations related to stress, ultimately influencing both physiological and psychological outcomes (Koelsch, 2015; Chanda & Levitin, 2013; Aalbers et al., 2017; Sakka & Juslin, 2018). Recently, research has increasingly focused on bodily responses, particularly interoceptive sensations, to better understand the relationship between music and emotion (Witte et al., 2019). A recent study by Putekin and colleagues (2024) utilized bodily sensation maps to visualize these responses (Nummenmaa, 2014; 2018) and found that music can evoke strong subjective bodily emotions that correlate with the emotional experiences that music elicits. These findings support models that highlight the importance of interoceptive sensations in the experience of emotions (Craig, 2002; Damasio and Carvalho, 2017). Additionally, Daikoku et al. (2024) identified specific body regions associated with emotional responses to music, revealing that positive emotions related to music are strongly felt in the chest area. In contrast, negative emotions such as anxiety and confusion are linked to sensations in the head region.

However, previous studies on music, emotions, and bodily responses have primarily focused on adults, taking into account various individual and cultural differences. While music is known to enhance psychological well-being across all age groups, there is currently a gap in research examining the relationship between music and these aspects across different generations, including school-age children and older adulthood, in a chronological context.

Therefore, we aim to clarify the differences in interoception and bodily sensations experienced while listening to music throughout the lifespan. We also seek to identify specific musical factors that contribute to promoting well-being. In particular, we examined bodily sensations during music listening and their connections to distinct emotional states. To achieve this, we conducted online experiments that included participants aged 10 to 80, focusing on body mapping and emotional assessments using music based on eight types of four-chord progressions. These chord progressions consider aspects of musical uncertainty, prediction errors, and temporal dynamics. The BRECVEMA framework proposed by Juslin (2013) and the research of Vuust and colleagues (2022) indicate that predictions related to music significantly influence its emotional impact. Furthermore, our previous studies have shown that these chord progressions are linked to various emotional components in music (Daikoku et al., 2024; Daikoku and Tanaka, 2024a, b; Tanaka and Daikoku, 2024.).

And we hypothesize that interoceptive and bodily sensations experienced while listening to music vary with age. Specifically, we suggest that these sensations gradually increase during adolescence as the body develops, peak during young adulthood, and subsequently decline in older age.

## Results

### 1. Relationship between age and Bodily sensations in each chord progression

The study first examined the relationship between bodily sensations and age groups for each chord progression. It analyzed bodily sensations separately for the whole body, the heart and abdomen (which are involved in interoceptive sensation), and the head, as previous research has noted frequent sensations in the head during sound listening.

We applied Spearman correlation tests to analyze the data, as the Shapiro–Wilk test for normality indicated significant violations of normality across all data (*p < .001*). Grand averages of the bodily sensation maps and the clicked positions for each chord progression were illustrated in Figures 2 and S2 in the supplemental information for each age group. All statistical analysis results, including descriptive statistics, have been made available in an external resource (https://osf.io/wzvqj/?view_only=4ecb8345b0c04917b9e219441a44def6).

**Figure 1.**
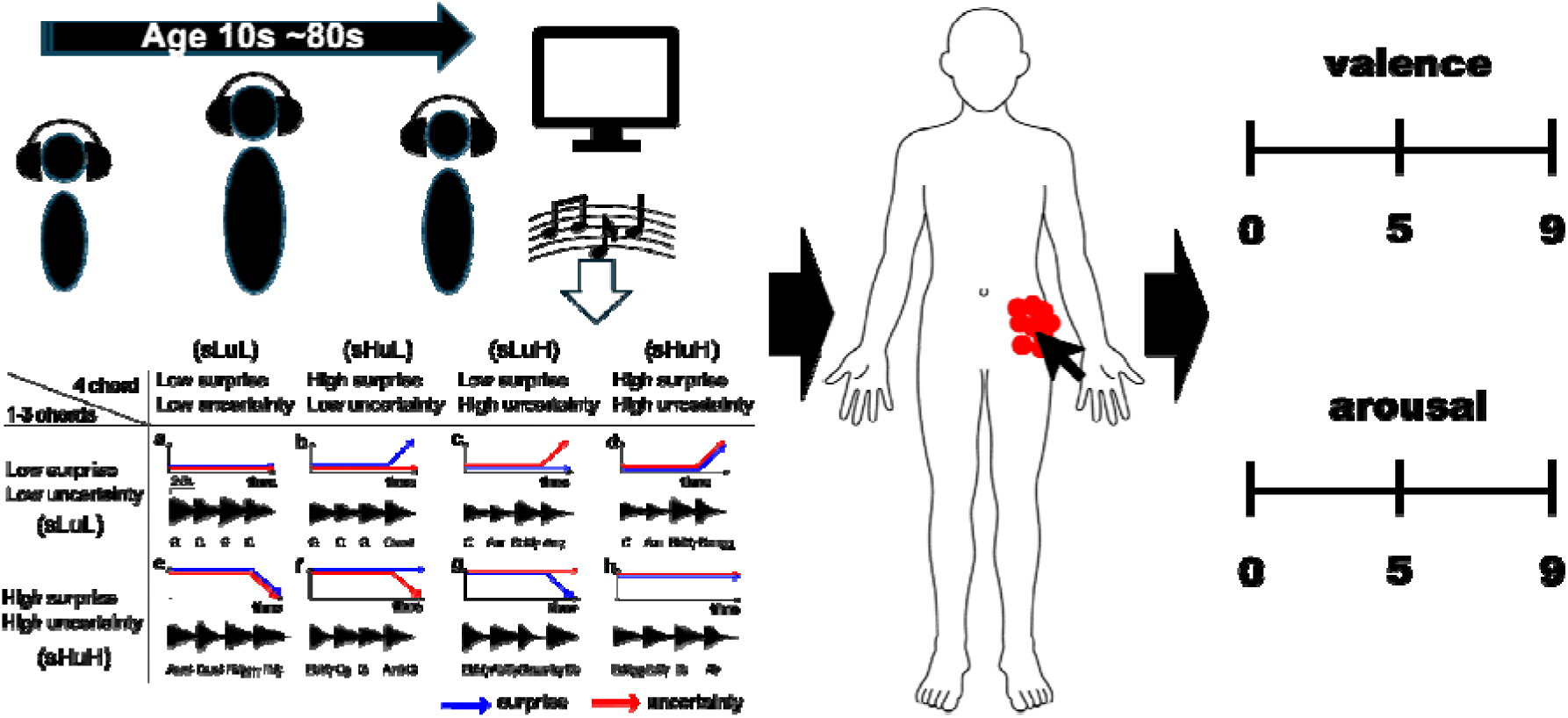
The experimental paradigm and sound stimuli. Below are 8 types of chord progressions. The blue and red arrows indicate surprise and uncertainty values, respectively.

**Figure 2.**
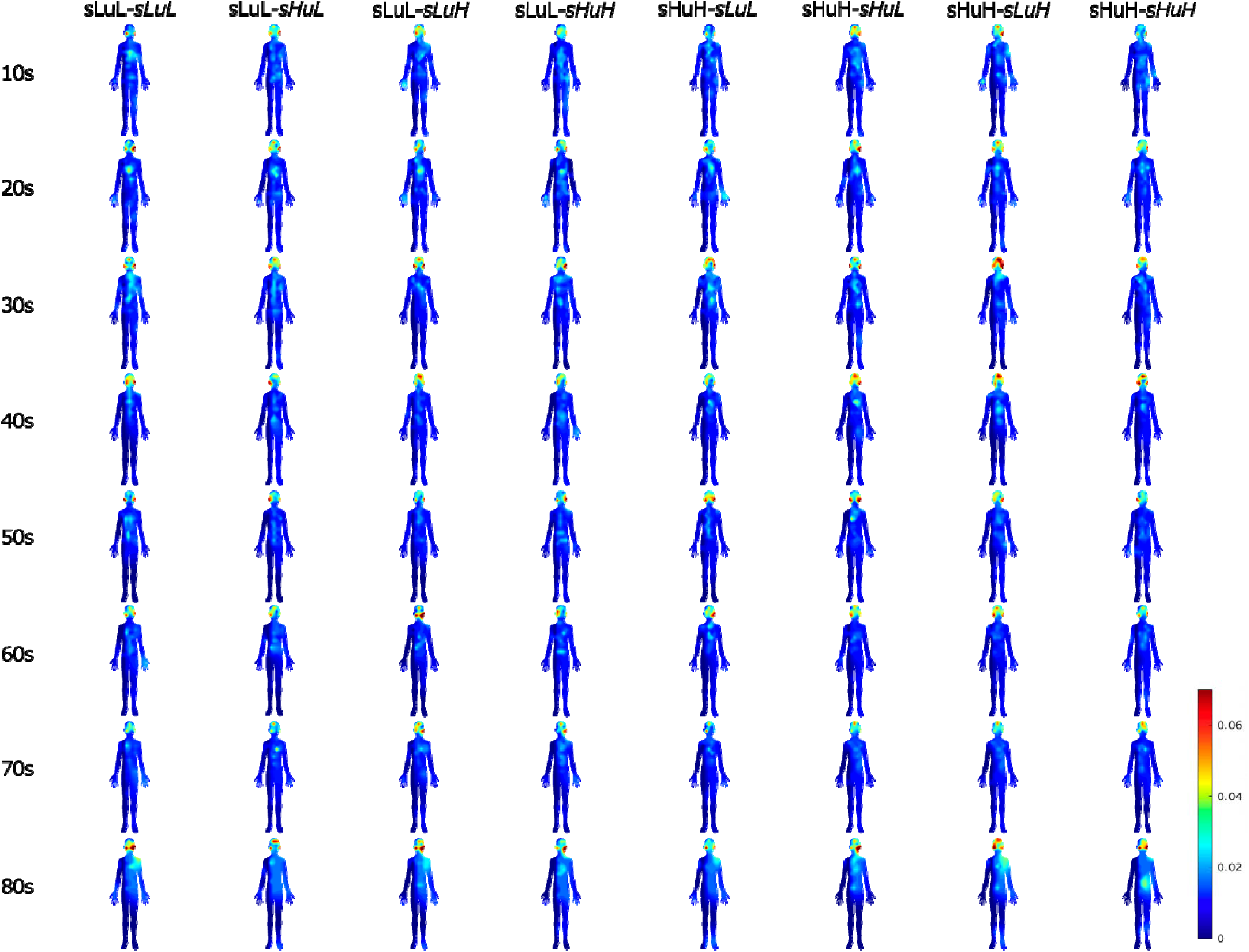
Body topography for chord progressions in the depression group. The blue-to-red gradients represent the number of clicks, ranging from few to many, respectively. Body responses peaked in the 30s for all chord chords and declined with increasing age.

The overall results revealed a correlation coefficient (*rho*) of –0.64 (*p < .001*) for the whole body (click SUM), –0.39 (*p < .001*) for the heart, and –0.40 (*p < .001*) for the abdomen, indicating a negative correlation between age and bodily sensation (regardless of location). The head showed no significant tendency, with a correlation of –0.19 (*p = .10*) (Figure 3).

**Figure 3.**
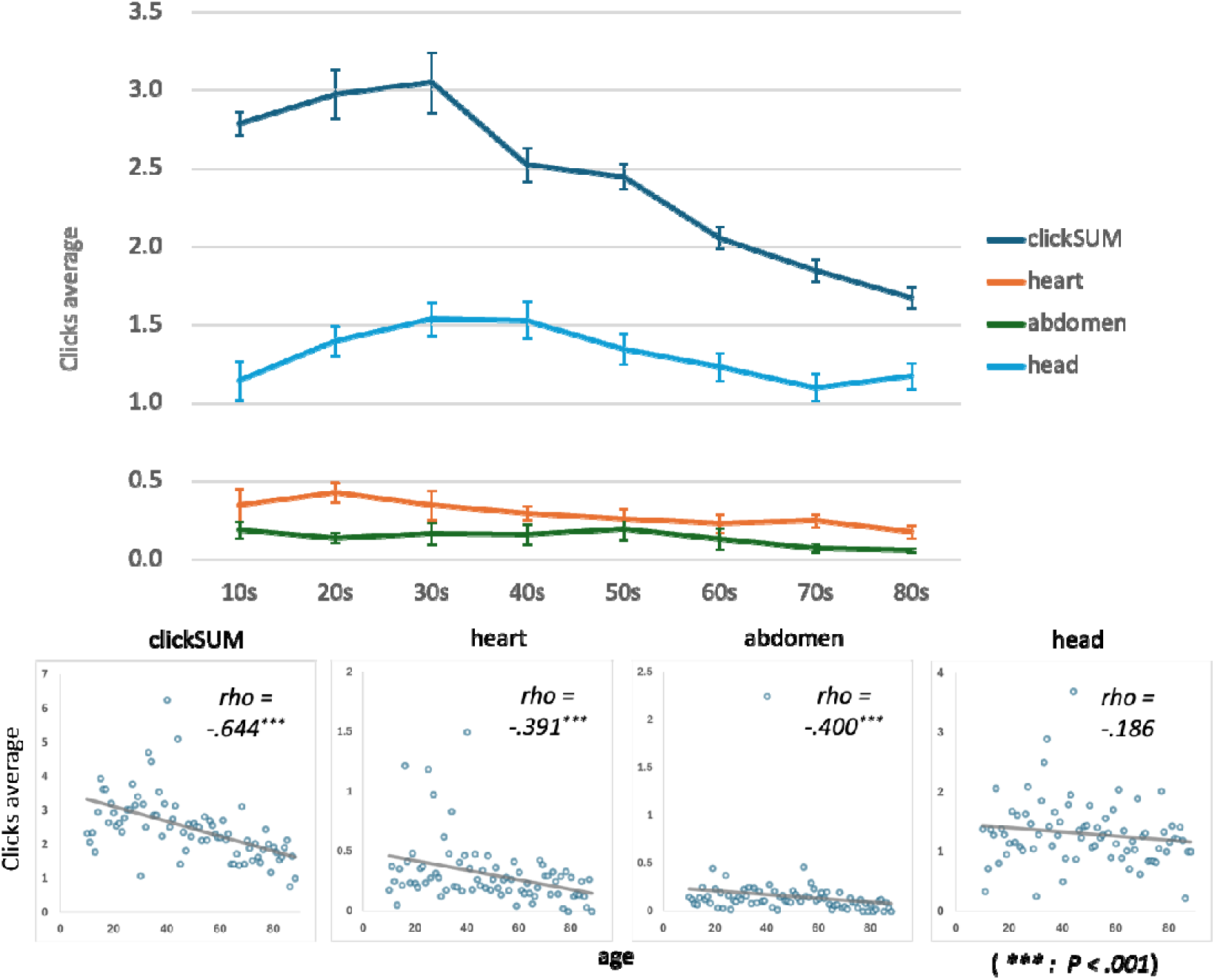
The average number of clicks by body map area by age. Negative correlations were shown for all body sensations (especially in areas related to interoception) with increasing age, except for the head.

Further examination of chord-specific results indicated a negative correlation between age and bodily sensations (excluding the head), particularly in chord progressions with low uncertainty and prediction error. For instance, the sLuL-sLuL progression showed a correlation of *rho* = –0.57 (*p < .001*) for the whole body, *rho =* –0.37 (*p < .001*) for the heart, *rho =* –0.27 (*p = .01*) for the abdomen, and *rho =* –0.11 (*p = .32*) for the head. Similarly, the sHuH-sLuL progression recorded *rho =* –0.53 (*p < .001*) for the whole body, *rho =* –0.31 (*p = .01*) for the heart, *rho =* –0.22 (*p = .05*) for the abdomen, and *rho =* –0.06 (*p = .63*) for the head. (Fig. 4).

**Figure 4.**
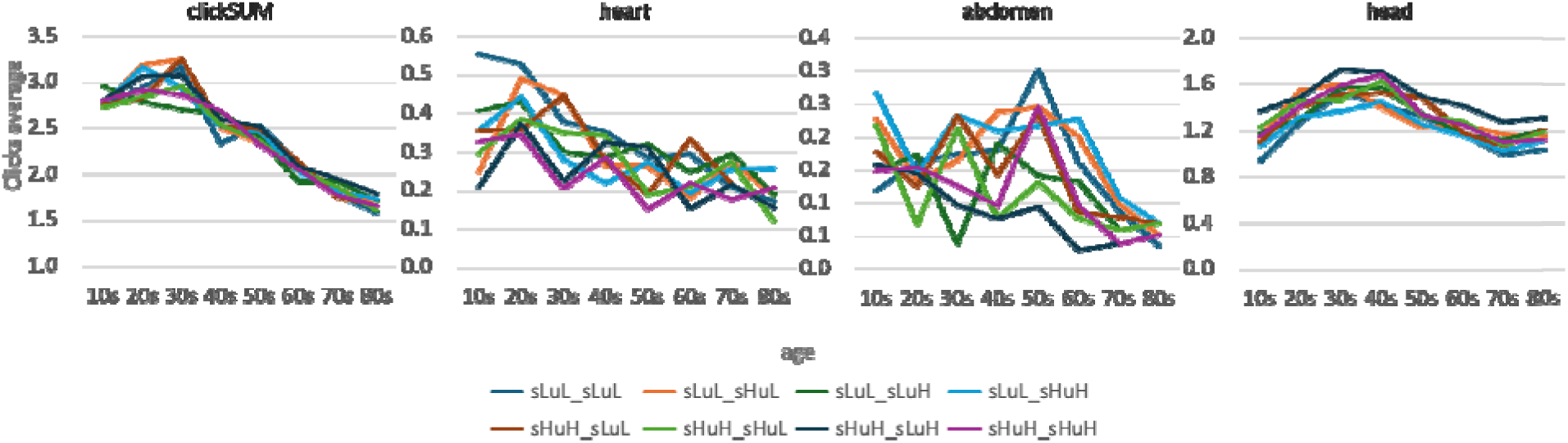
The average number of clicks per chord by age for body map area. Chord-specific results indicated a negative correlation between age and bodily sensations (excluding the head), particularly in chord progressions with low uncertainty and prediction error. And, for the body as a whole, all chords showed a decline in bodily sensations with increasing age, peaking in the 30s.

### 2. Comparing between age and each chord progressions in bodily sensations

We conducted a comparative analysis between age groups and each chord progression for different body parts based on the observed trends. The results of the paired test from the one-way analysis of variance revealed that, for the body as a whole, the following sequences were significantly higher in the individuals in the 30s age group compared to individuals in the 60s: sLuL-sLuL sequence (*w = –4.13, p = .02*), sLuL-sHuL sequence (*w = –5.31, p = .001*), sHuH-sLuL sequence (*w = –3.83, p = .04*), sHuH-sHuL sequence (*w = –4.59, p = .001*), sHuH-sLuH sequence (*w = –4.32, p = .01*). Additionally, the following sequences were significantly higher in the individuals in the 20s age group compared to individuals in the 60s:sLuL-sHuH sequence (*w = –5.18, p = .001*) and sHuH-sHuH sequence (*w = –4.91, p = .002*), This data suggests a general upward trend in these sequences, peaking in the young adulthood from 20s to 30s, as illustrated by the body map (Figure 2) and graphs (Figure 4). The only exception was the sLuL-sLuH sequence, which was also significantly higher in individuals in the 10s compared to individuals in the 60s (*w = –4.82, p = .001*), in individuals in the 30s compared to individuals in the 60s (*w = –5.30, p = .001*), and in individuals in the 20s compared to individuals in the 60s (*w = –4.32, p = .01*).

Regarding heart click counts, a peak was observed in individuals in the 20s for several chord sequences (Figure 2 & Figure 4). Specifically, the following sequences were significantly higher in individuals in the 20s compared to individuals in the 60s: sLuL-sHuL sequence (*w = –2.98, p = .01*), sLuL-sHuH sequence (*w = –2.34, p = .03*), and sHuH-sLuH sequence (*w = –4.04, p = .01*). Additionally, the sLuL-sLuL sequence (*w = –2.26, p = .03*) was significantly higher in individuals in the 20s compared to individuals in the 70s, while the sLuL-sLuH sequence (*w = –2.05, p = .03*) and the sHuH-sHuL sequence (*w = –3.14, p = .02*) were significantly higher in individuals in the 20s compared to individuals in the 50s.

Concerning the abdomen, there was a noticeable tendency towards a decrease in bodily sensation with increasing age. However, we did not identify clear directional changes related to specific chord progressions in the statistical analysis.

For the head, Figure 4 illustrates an inverted U-shape trend, with peaks observed in individuals in their 30s and 40s. Specifically, the sLuL-sLuL sequence was significantly higher in individuals in the 30s compared to individuals in the 10s (*w = 4.27, p = .002*) and the 70s (*w = –3.37, p = .03*). Similarly, the sLuL-sHuH sequence was significantly higher in individuals in the 30s compared to individuals in the 10s (*w = 2.94, p = .03*) and the 70s (*w = –2.98, p = .01*). Furthermore, the sHuH-sHuL sequence was significantly higher in individuals in individuals in the 40s compared to individuals in the 10s (*w = 3.01, p = .01*) and the 70s (*w = –0.74, p = .04*).

### 3. Relation between age and emotions

The results indicated a trend where arousal levels increased with age, showing an average increase in Likert scale score for all codes (*rho = .24, p = .04*) (Figure 5). In contrast, the valence ratings displayed a negative trend, though it was not statistically significant. A strong positive correlation was observed between the predictability features of the chords, specifically between sLuL-sLuL(*rho = .29, p = .01*) and sHuH-sLuL(*rho = .32, p = .003*), and arousal. No dominant trends were identified for valence.

**Figure 5.**
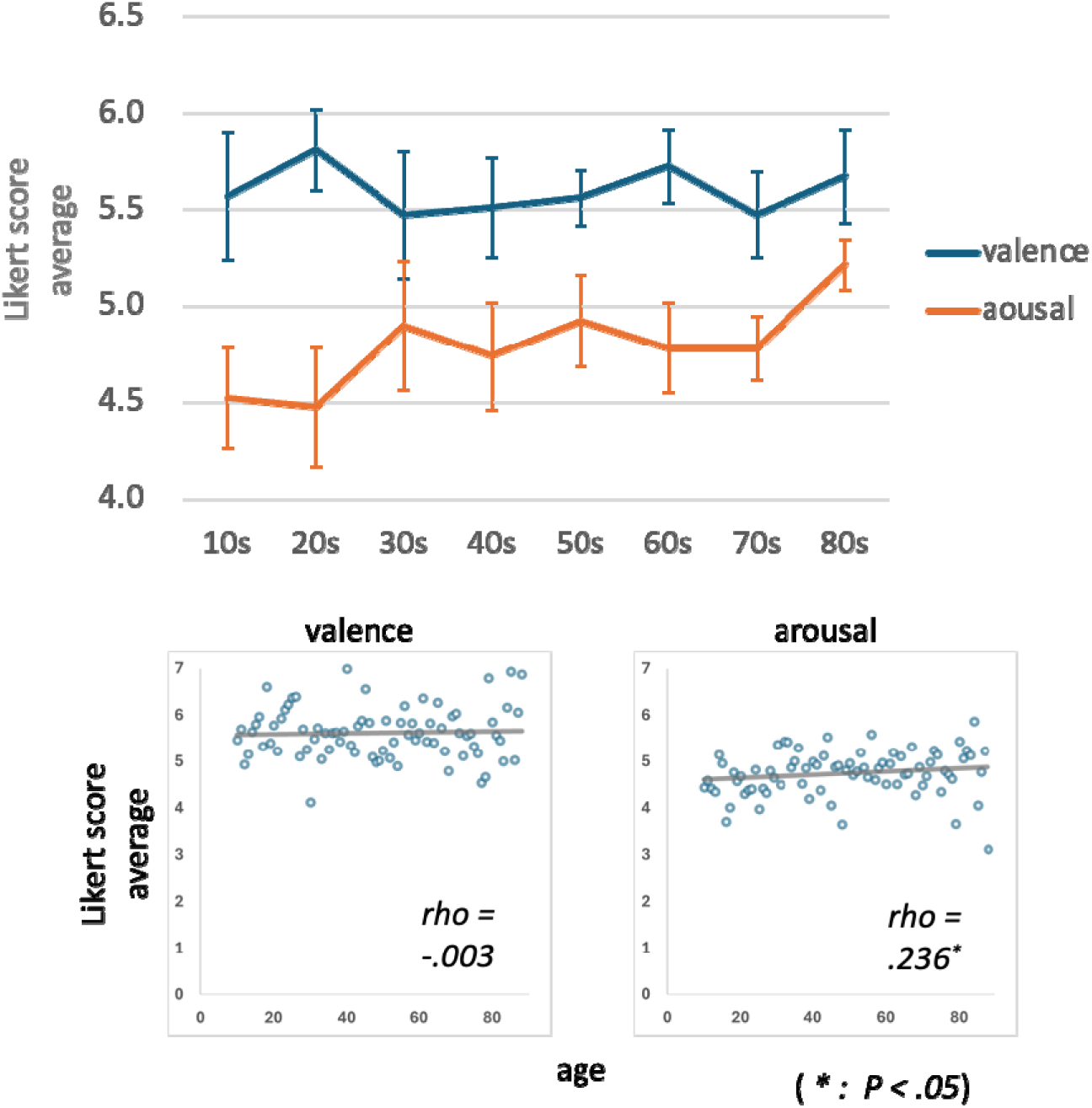
Valence and Arousal for all chord progressions by ages. For arousal, Likert scale scores for all codes increased on average, but no statistical advantage was shown for valence, although a negative trend was observed.

Regarding the relationship between age and emotional coding, valence showed variations in dominance among different age groups, but there was no clear trend over time. For arousal, significant differences were found across ages with sLuL-sLuL (χ*² = 17.08, p = .02,* ε*² = .02*) and sHuH-sLuL (χ*² = 17.41, p = .02,* ε*² = .02*). The code sLuL-sLuL was significantly more prevalent in individuals in their 80s compared to those in their 10s (*w = 4.40, p = .001*) and 20s (*w = 4.31, p = .003*) based on paired tests. Similarly, sHuH-sLuL was significantly higher in individuals in the 80s than in individuals both in the 10s (*w = 3.98, p = .01*) and in the 20s (*w = 4.43, p = .001*).

## Discussion

In this study, we aimed to clarify the differences in interoception and bodily sensations experienced while listening to music throughout the lifespan. We hypothesized that interoceptive and bodily sensations experienced while listening to music vary with age. We expected these sensations to gradually increase from school age as physical development progresses, peak during young adulthood, and then decline toward old age. Our results supported this hypothesis. Across all musical chords, we found a negative correlation between age and bodily sensations, regardless of their location. This trend was particularly evident for sounds with low uncertainty and low prediction error, although it was not observed for sensations localized in the head. Bodily sensations for all chords showed a decline after peaking in the 30s. Specifically, sensations in the chest peaked in the 20s and then decreased, except for sounds characterized by high uncertainty and prediction error at the beginning and low levels at the end, or consistently high levels throughout. In the abdomen, we observed a general decreasing trend, but no significant age-related differences were detected. In contrast, for head sensations, sounds where the beginning is low in both uncertainty and prediction error and the end is high or low in both exhibited an inverted U-shaped curve, peaking in the 30s.

Regarding emotions elicited by chords, regardless of the initial conditions, chords with both low uncertainty and low prediction error showed a strong positive correlation with arousal. This suggests that arousal increases with aging. However, for valence, while there appeared to be a negative trend between age and bodily sensations, no statistically significant differences were found.

When examining the emotions evoked by musical chords, regardless of their beginning, arousal showed a strong positive correlation with chords whose endings had low uncertainty and prediction error. Additionally, a positive correlation was observed for chords whose ending had high uncertainty and prediction error. These findings suggest that arousal tends to increase with aging. However, although no statistically significant differences were identified when considering valence, there appeared to be a negative trend between age and bodily sensations.

Humans inherently integrate top-down stimulus predictability with bottom-up interoceptive sensations (Seth et al., 2011; Seth and Critchley, 2013; Barrett and Simmons, 2015; Ondobaka et al., 2017; Quattrocki and Friston, 2014; Owens et al., 2018). Awareness of these bottom-up interoceptive sensations requires sensitivity to interoceptive signals transmitted through various sensors, such as mechanoreceptors and chemoreceptors. These sensors connect the peripheral and central nervous systems along the axes of the heart, respiratory system, and stomach (Berntson & Khalsa, 2021).

According to a review by Murphy and colleagues (2017), interoceptive ability, as assessed by a heartbeat discrimination task, has been shown to perform comparably in children (ages 6–11) and adults (Koch and Pollatos, 2014). Although research is limited, these abilities are thought to develop during childhood. However, studies indicate that interoceptive abilities, like exteroception and proprioception, decline with aging due to reduced sensitivity in the central nervous system and peripheral receptors (Khalsa et al., 2009).

Our findings indicate that, although bodily responses to auditory stimuli accompanied by predictions generally decrease with age after peaking during young adulthood, this effect is particularly pronounced for stimuli that are easier to predict and align with predictions—namely, those with low prediction error and low uncertainty. This trend aligns with previous research but highlights the heightened impact of predictability on bodily responses in this context aging.

Previous research on age-related differences in top-down stimulus predictability indicates that children, due to their limited past experiences, place a strong emphasis on the novelty of incoming stimuli and rely more on episodic memory when processing new information. In contrast, as the brain matures beyond young adulthood, individuals develop and maintain more refined predictive models. Older adulthood, in particular, are shown to rely significantly on their extensive knowledge base and long-term semantic memory representations acquired over a lifetime. This leads to a greater reliance on top-down predictions (Umanath & Marsh, 2014; Trapp et al., 2022; Shing et al., 2023).

In this study, we observed that arousal tends to increase with age. It was shown to be particularly pronounced at low prediction error and low uncertainty, the same chordal predictions as age-related reductions in physical response. However, it is difficult to definitively link this finding to differences in the relationship between prediction error or uncertainty based on our current results. Further research is needed to clarify these relationships. Nonetheless, the increase in arousal with age may be associated with enhanced brain activity, which is likely driven by top-down processing influenced by accumulated experience.

Also, we found no significant relationship between valence and the measured outcomes. However, our previous research using the same chord stimuli revealed a positive correlation between cardiac sensations and valence, though this was observed only in adults (Daikoku et al., 2024). Given the longitudinal study results from school age, adults, and older adulthood in this study—indicating a decline in cognitive valence with age, along with a reduction in bodily sensations—it is possible that bodily sensations are indeed related to valence.

The World Health Organization (WHO, 2000) defines well-being as “a state in which an individual realizes their own abilities, can cope with the normal stresses of life, works productively and fruitfully, and can contribute to their community.” The concept of well-being is recognized as multifaceted, and research commonly distinguishes between two dimensions: hedonia, which refers to the experience of positive emotions, and eudaimonia, which denotes optimal functioning (Hallam and Himonides, 2022). Therefore, this cross-sectional study is expected to shed significant light on research aimed at promoting multifaceted well-being by elucidating differences in bodily sensory responses and emotions induced by music, which are likely associated with developmental changes in physiological functions and the brain.

This study has several limitations. First, the experiments were conducted online, which may have compromised the experimental control compared to a laboratory setting. Second, we did not consider the potential influence of factors such as antihypertensive medications, which are commonly prescribed in older adulthood and may affect bodily responses (Ulus and Aisenberg-Shafran, 2022). Nevertheless, since the study aims to explore music that promotes well-being, it is also important to understand the effects of music in everyday life and lifestyle contexts, rather than solely in controlled laboratory environments.

In summary, this study found that listening to specific chord stimuli may enhance arousal due to the influence of top-down stimulus prediction, which is shaped by accumulated experiences as we age. In contrast, bottom-up bodily sensations tend to diminish as adults grow older. Although no definitive relationships were established in this current study, our previous research involving adult participants indicated a link between cardiac regions and bodily responses. These findings contribute to our understanding of how music affects us physically and psychologically, and they may shed light on identifying music that promotes well-being while taking into account generational differences in physical and mental characteristics.

## STAR Methods

### Participants

The present study involved conducting bodily sensation mapping tests along with emotional judgments for each of the eight types of 4-chord progressions (for details, see Daikoku and Tanaka, 2024a). A total of 784 Japanese participants, aged between 10 and 80 years (mean age ± SD = 47.42 ± 21.34, including 373 females), took part in the research. All participants had no history of neurological or audiological disorders and did not possess absolute pitch. The experiment was conducted in accordance with the guidelines of the Declaration of Helsinki and received approval from the Ethics Committee of The University of Tokyo (Approval No. UT IST RE 230601). All participants provided informed consent and conducted the experiments on a PC.

### Materials

The experimental paradigm was developed using the Gorilla Experiment Builder, a cloud-based research platform that allows for the online deployment of behavioral experiments. Each participant listened to eight different four-chord progressions, with each chord lasting 500 milliseconds. The audio was played at a sampling rate of 44.1 kHz and in 32-bit format, using General MIDI’s Electric Piano 1 sound. To ensure consistent loudness perception among all participants, the amplitudes of the chords were adjusted based on equal loudness contours.

A statistical learning model (Daikoku, Minaotya et al., 2023) was employed to estimate the surprise and uncertainty associated with each chord. This analysis was based on data from the McGill Billboard corpus (Burgoyne, Wild, and Fujinaga, 2011), which consists of 890 pop songs. Shannon information content and entropy were calculated based on transition probabilities (Shannon, 1948). Transition probability is often interpreted as the amount of information present; a lower amount of information (indicating a higher transition probability) results in greater predictability and fewer surprises, while a higher amount of information (indicating a lower transition probability) leads to less predictability and more surprises. Entropy measures the listener’s uncertainty when predicting the next chord based on the previous one, while the amount of information indicates the level of surprise experienced upon hearing a chord. Using this framework, we developed 92 distinct chord progressions, which we classified into eight different types.

The eight types of chord progressions can be categorized based on the uncertainty and surprise of their chords. Four of these types begin with three chords that exhibit low uncertainty and surprise, while the other four start with three chords that feature high uncertainty and surprise. In each set of four progressions, the fourth chord varies in both uncertainty and surprise. The types are as follows:

1) sLuL-sLuL sequence (Figure 1.a): the 1st to 3rd chords have low surprise and uncertainty and the 4th chord also has low surprise and uncertainty,
2) sLuL-sHuL sequence (Figure 1.b): the 1st to 3rd chords have low surprise and uncertainty, while the 4th chord has high surprise and low uncertainty,
3) sLuL-sLuH sequence (Figure 1.c): the 1st to 3rd chords have low surprise and uncertainty, while the 4th chord has low surprise and high uncertainty,
4) sLuL-sHuH sequence (Figure 1.d): the 1st to 3rd chords have low surprise and uncertainty, while the 4th chord has high surprise and uncertainty,
5) sHuH-sLuL sequence (Figure 1.e): the 1st to 3rd chords have high surprise and uncertainty, while the 4th chord has low surprise and uncertainty,
6) sHuH-sHuL sequence (Figure 1.f): the 1st to 3rd chords have high surprise and uncertainty, while the 4th chord has high surprise and low uncertainty,
7) sHuH-sLuH sequence (Figure 1.g): the 1st to 3rd chords have high surprise and uncertainty, while the 4th chord has low surprise and high uncertainty,
8) sHuH-sHuH sequence (Figure 1.h): the 1st to 3rd chords have high surprise and uncertainty, and the 4th chord also has high surprise and uncertainty.

A variety of chord progressions were generated for each of these eight types: sLuL-sLuL (14 progressions), sLuL-sHuL (14), sLuL-sLuH (12), sLuL-sHuH (8), sHuH-sLuL (18), sHuH-sHuL (14), sHuH-sLuH (3), and sHuH-sHuH (9). The thresholds for high and low uncertainty and surprise were determined based on the top 20% and bottom 20% of all data points.

### Procedures

Participants were presented with eight different chord progressions in a random order. After each session, they were asked to respond within three seconds by clicking on a body image displayed on the screen, indicating where they felt sensations from the chords. Participants could click multiple times if they wished (see Figure S1 in the supplementary material for details). Additionally, each participant completed a survey for emotional judgment. They were asked to rate each type of chord progression based on valence and arousal. Each rating was made using a nine-point Likert scale, where the midpoint, number 5, represented a neutral response.

### Quantification and statistical analysis

Using the coordinate data of x and y from the body mapping test, we extracted the total number of clicks at two interoceptive positions: the cardiac and abdominal areas for each participant. The raw x and y coordinate data (see Figure S2 in the supplementary material) were downsampled by a factor of 40. The body image template used measured 871 pixels in width and 1920 pixels in height. The body topography figures (Figure 2) were generated using Matlab (2022b) by interpolating the x and y coordinates in a mesh grid format, employing a color map to represent the neighboring points.

We performed the Shapiro–Wilk test for normality on the different click positions, the total number of clicks for each participant, and the total clicks at the cardiac, abdominal, and head positions, as well as on the valence and arousal scores from the nine-point dimensional judgments. Depending on the normality test results, we applied either parametric or non-parametric (Spearman) correlation analyses to examine the relationships between the average age, click counts at the cardiac, abdominal, and head positions, total click counts, and emotion scores. Additionally, we conducted a non-parametric (Kruskal-Wallis) One-Way analysis of variance (ANOVA) to compare age differences among different types of chord progressions in relation to body sensations and emotions. Statistical analyses were carried out using jamovi Version 1.2 (The jamovi project, 2021). We set the threshold for statistical significance at p < .05 and employed a false discovery rate (FDR) for post-hoc analysis and multiple comparisons.

## Data Availability

All anonymized raw data files, stimuli used in this study and the results of statistical analysis have been deposited to an external source (https://osf.io/wzvqj/?view_only=4ecb8345b0c04917b9e219441a44def6). The other data are shown in supplementary data.

## Acknowledgements

This research was supported by the Japan Society for the Promotion of Science (JSPS), Fostering Joint International Research (22KK0157), Japan. The funding sources had no role in the decision to publish or prepare the manuscript.

## Author Contributions Statement

M.T. and T.D. conceived the experimental paradigm and method of data analysis. M.T. collected the data, analysed the data, and wrote the draft of the manuscript and figure. M.T. and T.D. edited and finalized the manuscript.

