## Supplementary for "Lifetime changes in Body map based on music prediction"

Figure S1. Body image and example of the clicks to the position in the body.

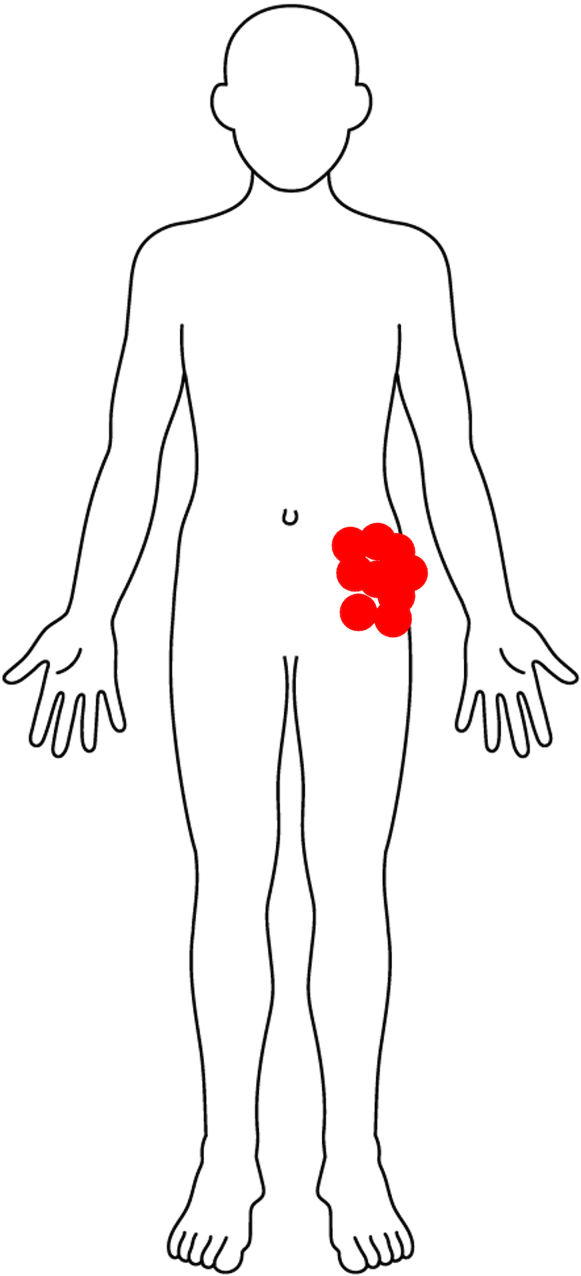

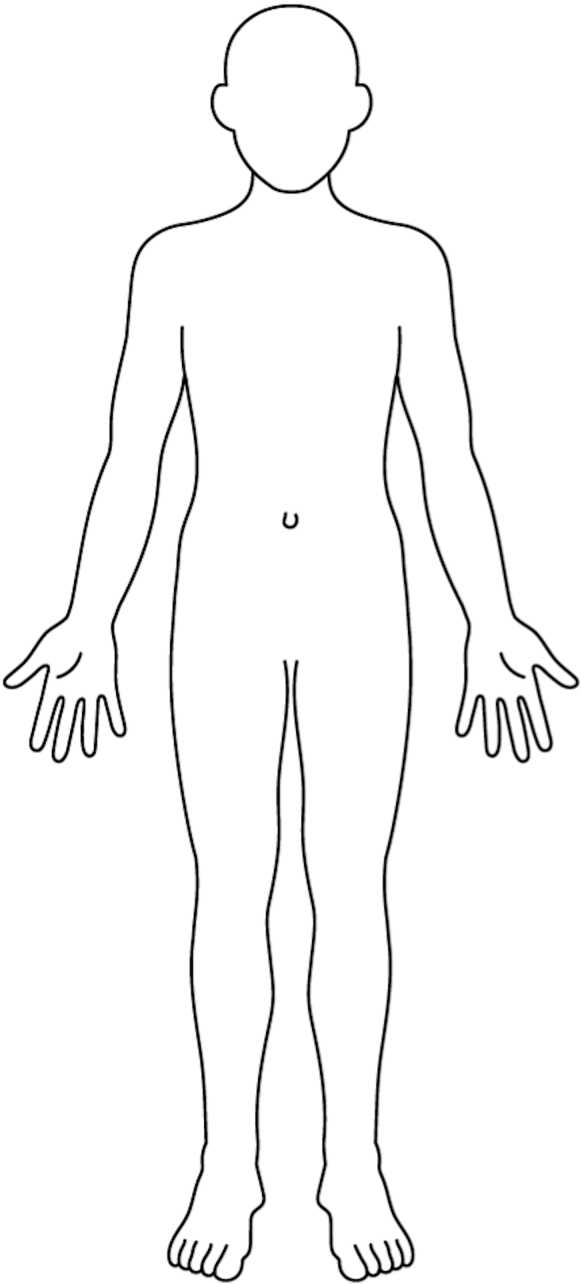

Figure.S2. Clicked body positions in each chord progression for each age group

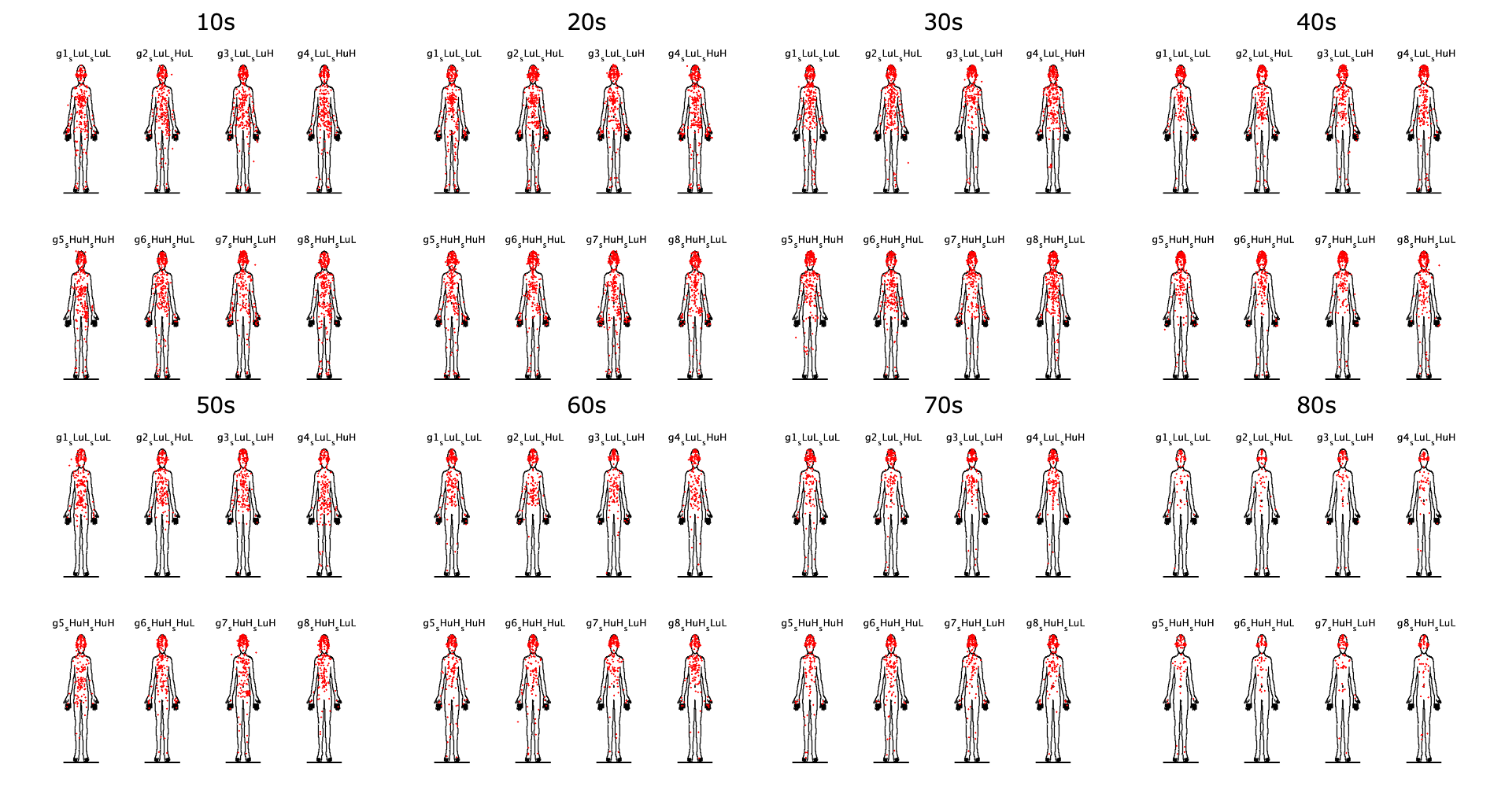
